## Supplementary Figures 1-3 and Tables 2-7 for "High-throughput identification of prefusion-stabilizing mutations in SARS-CoV-2 spike"

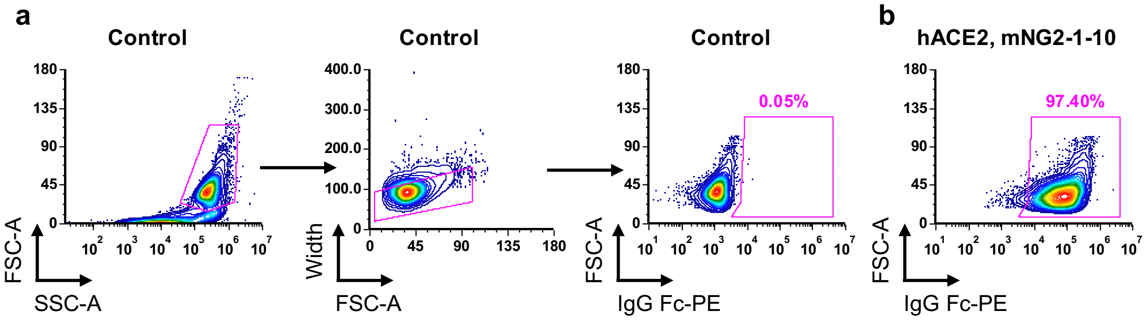


**Supplementary Fig. 1 | Validation of hACE2, mNG2$_{1-10}$-expressing cell line. a,b,** Flow cytometry plots showing (**a**) untransfected control and gating strategy in this verification experiment, and (**b**) surface expression of hACE2 in HEK293T landing pad cells.

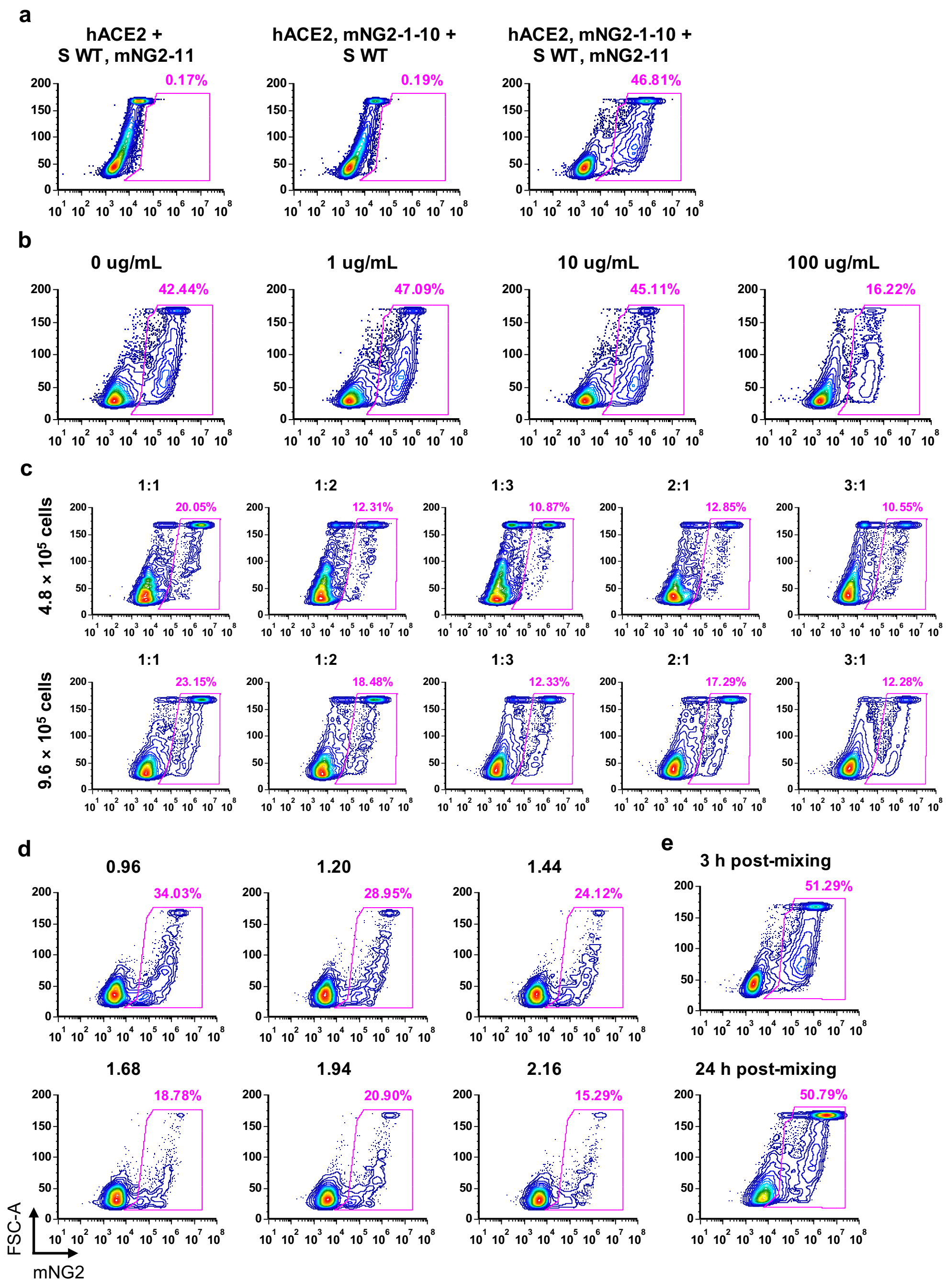


**Supplementary Fig. 2 | Validation and optimization of fusion assay. a,** Flow cytometry plots of fusion assay between hACE2- and S-expressing cells showing detectable green fluorescence when both mNG2$_{1\text{-}10}$ and mNG2$_{11}$ were expressed, and negligible fluorescent background signal when either mNG2$_{1\text{-}10}$ or mNG2$_{11}$ was expressed. **b,** Addition of CC40.8, a neutralizing antibody targeting the stem helix, to S, mNG2$_{11}$-expressing cells one hour prior to mixing with hACE2,

$mNG2_{1\text{-}10}$-expressing cells decreased fusion events. Antibody concentration is indicated above each plot. **c,** Optimization of the ratio between hACE2, $mNG2_{1\text{-}10}$- and S, $mNG2_{11}$-expressing cells for fusion assay. Total cell number in co-culture is indicated, and the ratio of hACE2, $mNG2_{1\text{-}10}$- to S, $mNG2_{11}$-expressing cells is shown above each plot. **d,** Optimization of total cell numbers of hACE2, $mNG2_{1\text{-}10}$- and S, $mNG2_{11}$-expressing cells for fusion assay. Total cell number, in millions, of hACE2, $mNG2_{1\text{-}10}$- and S, $mNG2_{11}$-expressing cells in co-culture is shown above each plot. Equal numbers of hACE2, $mNG2_{1\text{-}10}$- and S, $mNG2_{11}$-expressing cells were co-cultured for 3 hours. **e,** Optimization of time point for fusion sorting. Equal numbers of hACE2, $mNG2_{1\text{-}10}$- and S, $mNG2_{11}$-expressing cells totaling $5.0 \times 10^5$ cells/mL were mixed and analyzed via flow cytometry at the indicated time point.

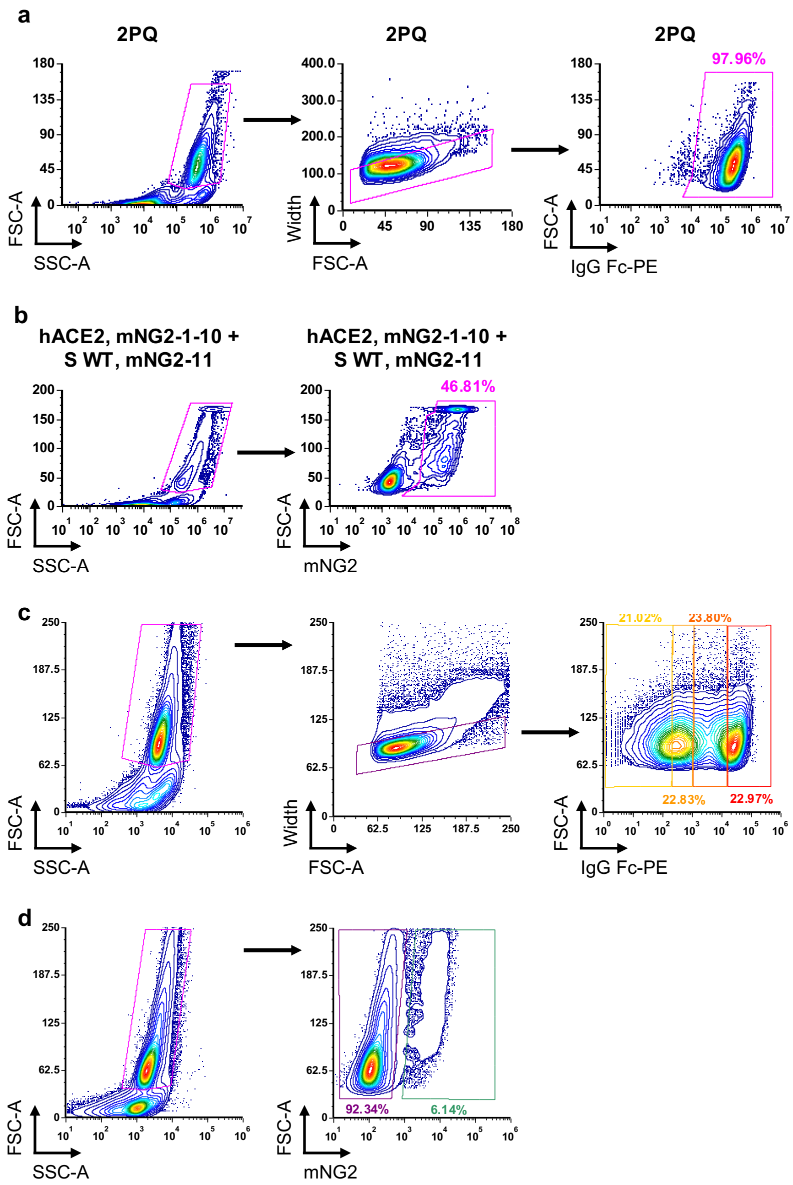


**Supplementary Fig. 3 | Gating strategies for flow cytometry and fluorescence activated cell sorting (FACS). a,** Gating strategy used for flow cytometry to assess surface expression of S via PE fluorescence. **b,** Gating strategy used for flow cytometry to determine fusion of hACE2- and S-expressing cells via mNG2 fluorescence. **c,** Gating strategy used for FACS to sort the DMS library of S-expressing cells based on levels of PE fluorescence. **d,** Gating strategy used for FACS to sort the co-culture of the DMS library of S-expressing cells and hACE2-expressing cells at 3 hours post-mixing, based on presence or absence of mNG2 fluorescence.

**Supplementary Table 2.** Primers used for PCR-based site directed mutagenesis (QuikChange) to generate mutations. The plasmid backbone used was attB-S-mNG2-11.

| **Primer** | **Sequence (5′ to 3′)** |
|---|---|
| T961F-F | GCACTGAACTTCCTGGTCAAGCAGCTGTCC |
| T961F-R | CAGCTGCTTGACCAGGAAGTTCAGTGCCTGGGC |
| K986P-F | CTGAGCAGACTGGACCCGGTGGAAGCCGAGGTGCAG |
| K986P-R | CTGCACCTCGGCTTCCACCGGGTCCAGTCTGCTCAG |
| V987P-F | CTGAGCAGACTGGACAAGCCGGAAGCCGAGGTGCAG |
| V987P-R | CTGCACCTCGGCTTCCGGCTTGTCCAGTCTGCTCAG |
| D994E-F | CGAGGTGCAGATCGAGAGACTGATCACCGGAAG |
| D994E-R | CTTCCGGTGATCAGTCTCTCGATCTGCACCTCGGCT |
| D994Q-F | GCCGAGGTGCAGATCCAGAGACTGATCACCGGAAG |
| D994Q-R | CTTCCGGTGATCAGTCTCTGGATCTGCACCTCGGCT |
| Q1005R-F | GGCTGCAGTCCCTGCGGACCTACGTTACCCAG |
| Q1005R-R | CTGGGTAACGTAGGTCCGCAGGGACTGCAGCC |
| K986P/V987P-F | CTGAGCAGACTGGACCCGCCGGAAGCCGAGGTGCAG |
| K986P/V987P-R | CTGCACCTCGGCTTCCGGCGGGTCCAGTCTGCTCAG |

32 **Supplementary Table 3.** Cassette primers to generate the DMS library.

| Primer | Sequence (5′ to 3′) |
|---|---|
| Cassette1_1 | GCCCTGCTGGCCGGCACAATCNNKAGTGGTTGGACATTTGGAGCTGGCGCCGCTCTGCAG |
| Cassette1_2 | GCCCTGCTGGCCGGCACAATCACCNNKGGCTGGACCTTTGGAGCTGGCGCCGCTCTGCAG |
| Cassette1_3 | GCCCTGCTGGCCGGCACAATCACCAGCNNKTGGACATTCGGAGCTGGCGCCGCTCTGCAG |
| Cassette1_4 | GCCCTGCTGGCCGGCACAATCACAAGCGGTNNKACCTTTGGAGCTGGCGCCGCTCTGCAG |
| Cassette1_5 | GCCCTGCTGGCCGGCACAATCACAAGTGGCTGGNNKTTCGGAGCTGGCGCCGCTCTGCAG |
| Cassette1_6 | GCCCTGCTGGCCGGCACAATCACCAGCGGTTGGACCNNKGGTGCTGGCGCCGCTCTGCAG |
| Cassette1_7 | GCCCTGCTGGCCGGCACAATCACCAGCGGTTGGACATTTNNKGCAGGCGCCGCTCTGCAG |
| Cassette1_8 | GCCCTGCTGGCCGGCACAATCACCAGTGGTTGGACCTTCGGANNKGGCGCCGCTCTGCAG |
| Cassette2_1 | AGCGGCTGGACATTTGGAGCTNNKGCAGCACTGCAGATCCCCTTTGCTATGCAGATGGCC |
| Cassette2_2 | AGCGGCTGGACATTTGGAGCTGGTNNKGCTCTCCAGATCCCCTTTGCTATGCAGATGGCC |
| Cassette2_3 | AGCGGCTGGACATTTGGAGCTGGTGCCNNKCTGCAAATCCCCTTTGCTATGCAGATGGCC |
| Cassette2_4 | AGCGGCTGGACATTTGGAGCTGGCGCAGCTNNKCAAATCCCCTTTGCTATGCAGATGGCC |
| Cassette2_5 | AGCGGCTGGACATTTGGAGCTGGCGCCGCACTCNNKATCCCCTTTGCTATGCAGATGGCC |
| Cassette2_6 | AGCGGCTGGACATTTGGAGCTGGTGCAGCACTCCAANNKCCCTTTGCTATGCAGATGGCC |
| Cassette2_7 | AGCGGCTGGACATTTGGAGCTGGTGCAGCTCTGCAAATANNKTTTGCTATGCAGATGGCC |
| Cassette2_8 | AGCGGCTGGACATTTGGAGCTGGTGCCGCACTCCAGATACCCNNKGCTATGCAGATGGCC |
| Cassette3_1 | GCCGCTCTGCAGATCCCCTTTNNKATGCAAATGGCATACCGGTTCAACGGCATCGGAGTG |
| Cassette3_2 | GCCGCTCTGCAGATCCCCTTTGCANNKCAAATGGCCTATCGGTTCAACGGCATCGGAGTG |
| Cassette3_3 | GCCGCTCTGCAGATCCCCTTTGCAATGNNKATGGCATATCGATTCAACGGCATCGGAGTG |
| Cassette3_4 | GCCGCTCTGCAGATCCCCTTTGCTATGCAGNNKGCATATCGGTTCAACGGCATCGGAGTG |
| Cassette3_5 | GCCGCTCTGCAGATCCCCTTTGCAATGCAAATGNNKTACCGATTTAACGGCATCGGAGTG |
| Cassette3_6 | GCCGCTCTGCAGATCCCCTTTGCTATGCAAATGGCCNNKCGATTCAACGGCATCGGAGTG |
| Cassette3_7 | GCCGCTCTGCAGATCCCCTTTGCAATGCAGATGGCCTATNNKTTTAACGGCATCGGAGTG |

| | |
|---|---|
| Cassette3_8 | GCCGCTCTGCAGATCCCCTTTGCTATGCAGATGGCATACCGANNKAA CGGCATCGGAGTG |
| Cassette4_1 | ATGCAGATGGCCTACCGGTTCNNKGGTATAGGAGTGACCCAGAATGT GCTGTACGAGAAC |
| Cassette4_2 | ATGCAGATGGCCTACCGGTTCAATNNKATCGGTGTGACCCAGAATGT GCTGTACGAGAAC |
| Cassette4_3 | ATGCAGATGGCCTACCGGTTCAACGGTNNKGGTGTAACCCAGAATGT GCTGTACGAGAAC |
| Cassette4_4 | ATGCAGATGGCCTACCGGTTCAATGGCATANNKGTAACCCAGAATGT GCTGTACGAGAAC |
| Cassette4_5 | ATGCAGATGGCCTACCGGTTCAACGGTATCGGANNKACACAGAATGT GCTGTACGAGAAC |
| Cassette4_6 | ATGCAGATGGCCTACCGGTTCAACGGCATAGGTGTGNNKCAGAATGT GCTGTACGAGAAC |
| Cassette4_7 | ATGCAGATGGCCTACCGGTTCAATGGTATCGGAGTAACCNNKAATGT GCTGTACGAGAAC |
| Cassette4_8 | ATGCAGATGGCCTACCGGTTCAACGGCATCGGTGTAACACAGNNKGT GCTGTACGAGAAC |
| Cassette5_1 | GGCATCGGAGTGACCCAGAATNNKCTCTATGAGAACCAGAAGCTGAT CGCCAACCAGTTC |
| Cassette5_2 | GGCATCGGAGTGACCCAGAATGTANNKTACGAAAACCAGAAGCTGAT CGCCAACCAGTTC |
| Cassette5_3 | GGCATCGGAGTGACCCAGAATGTACTGNNKGAGAATCAGAAGCTGAT CGCCAACCAGTTC |
| Cassette5_4 | GGCATCGGAGTGACCCAGAATGTGCTCTACNNKAATCAGAAGCTGAT CGCCAACCAGTTC |
| Cassette5_5 | GGCATCGGAGTGACCCAGAATGTGCTGTATGAANNKCAGAAGCTGAT CGCCAACCAGTTC |
| Cassette5_6 | GGCATCGGAGTGACCCAGAATGTACTCTATGAAAATNNKAAGCTGATC GCCAACCAGTTC |
| Cassette5_7 | GGCATCGGAGTGACCCAGAATGTACTCTACGAGAACCAANNKCTGAT CGCCAACCAGTTC |
| Cassette5_8 | GGCATCGGAGTGACCCAGAATGTACTGTATGAGAACCAAAAGNNKAT CGCCAACCAGTTC |
| Cassette6_1 | CTGTACGAGAACCAGAAGCTGNNKGCAAATCAGTTCAACAGCGCCAT CGGCAAGATCCAG |
| Cassette6_2 | CTGTACGAGAACCAGAAGCTGATANNKAACCAATTCAACAGCGCCATC GGCAAGATCCAG |
| Cassette6_3 | CTGTACGAGAACCAGAAGCTGATCGCANNKCAATTTAACAGCGCCATC GGCAAGATCCAG |
| Cassette6_4 | CTGTACGAGAACCAGAAGCTGATAGCCAATNNKTTTAACAGCGCCATC GGCAAGATCCAG |
| Cassette6_5 | CTGTACGAGAACCAGAAGCTGATCGCAAACCAGNNKAATAGCGCCAT CGGCAAGATCCAG |
| Cassette6_6 | CTGTACGAGAACCAGAAGCTGATCGCCAATCAATTCNNKAGCGCCAT CGGCAAGATCCAG |
| Cassette6_7 | CTGTACGAGAACCAGAAGCTGATAGCAAACCAGTTTAACNNKGCCATC GGCAAGATCCAG |
| Cassette6_8 | CTGTACGAGAACCAGAAGCTGATCGCCAACCAATTTAATAGCNNKATC GGCAAGATCCAG |

| | |
|---|---|
| Cassette7_1 | GCCAACCAGTTCAACAGCGCCNNKGGTAAAATCCAGGACAGCCTGAG CAGCACAGCAAGC |
| Cassette7_2 | GCCAACCAGTTCAACAGCGCCATANNKAAGATACAGGACAGCCTGAG CAGCACAGCAAGC |
| Cassette7_3 | GCCAACCAGTTCAACAGCGCCATCGGTNNKATACAAGACAGCCTGAG CAGCACAGCAAGC |
| Cassette7_4 | GCCAACCAGTTCAACAGCGCCATAGGCAAANNKCAAGACAGCCTGAG CAGCACAGCAAGC |
| Cassette7_5 | GCCAACCAGTTCAACAGCGCCATCGGTAAGATCNNKGATAGCCTGAG CAGCACAGCAAGC |
| Cassette7_6 | GCCAACCAGTTCAACAGCGCCATCGGCAAAATACAGNNKAGCCTGAG CAGCACAGCAAGC |
| Cassette7_7 | GCCAACCAGTTCAACAGCGCCATAGGTAAGATCCAAGACNNKCTGAG CAGCACAGCAAGC |
| Cassette7_8 | GCCAACCAGTTCAACAGCGCCATCGGCAAGATACAAGATAGCNNKAG CAGCACAGCAAGC |
| Cassette8_1 | GGCAAGATCCAGGACAGCCTGNNKAGTACCGCAAGCGCCCTGGGAA AGCTGCAGGACGTG |
| Cassette8_2 | GGCAAGATCCAGGACAGCCTGAGTNNKACAGCCAGCGCCCTGGGAA AGCTGCAGGACGTG |
| Cassette8_3 | GGCAAGATCCAGGACAGCCTGAGCAGCNNKGCCAGTGCCCTGGGAA AGCTGCAGGACGTG |
| Cassette8_4 | GGCAAGATCCAGGACAGCCTGAGCAGCACCNNKAGCGCACTGGGAA AGCTGCAGGACGTG |
| Cassette8_5 | GGCAAGATCCAGGACAGCCTGAGCAGTACAGCANNKGCACTGGGAAA GCTGCAGGACGTG |
| Cassette8_6 | GGCAAGATCCAGGACAGCCTGAGTAGCACCGCAAGTNNKCTGGGAAA GCTGCAGGACGTG |
| Cassette8_7 | GGCAAGATCCAGGACAGCCTGAGTAGTACCGCCAGTGCCNNKGGAAA GCTGCAGGACGTG |
| Cassette8_8 | GGCAAGATCCAGGACAGCCTGAGTAGTACAGCAAGTGCCCTGNNKAA GCTGCAGGACGTG |
| Cassette9_1 | AGCACAGCAAGCGCCCTGGGANNKCTCCAAGACGTGGTCAACCAGAA TGCCCAGGCACTG |
| Cassette9_2 | AGCACAGCAAGCGCCCTGGGAAAANNKCAGGATGTGGTCAACCAGAA TGCCCAGGCACTG |
| Cassette9_3 | AGCACAGCAAGCGCCCTGGGAAAGCTCNNKGATGTAGTCAACCAGAA TGCCCAGGCACTG |
| Cassette9_4 | AGCACAGCAAGCGCCCTGGGAAAACTGCAANNKGTAGTCAACCAGAA TGCCCAGGCACTG |
| Cassette9_5 | AGCACAGCAAGCGCCCTGGGAAAGCTCCAGGACNNKGTAAACCAGAA TGCCCAGGCACTG |
| Cassette9_6 | AGCACAGCAAGCGCCCTGGGAAAGCTGCAAGATGTGNNKAACCAGAA TGCCCAGGCACTG |
| Cassette9_7 | AGCACAGCAAGCGCCCTGGGAAAACTCCAGGACGTAGTCNNKCAGAA TGCCCAGGCACTG |
| Cassette9_8 | AGCACAGCAAGCGCCCTGGGAAAGCTGCAAGACGTAGTAAACNNKAA TGCCCAGGCACTG |
| Cassette10_1 | CTGCAGGACGTGGTCAACCAGNNKGCACAAGCACTGAACACCCTGGT CAAGCAGCTGTCC |

| | |
|---|---|
| Cassette10_2 | CTGCAGGACGTGGTCAACCAGAACNNKCAGGCCCTGAACACCCTGGT CAAGCAGCTGTCC |
| Cassette10_3 | CTGCAGGACGTGGTCAACCAGAACGCCNNKGCACTCAACACCCTGGT CAAGCAGCTGTCC |
| Cassette10_4 | CTGCAGGACGTGGTCAACCAGAATGCACAGNNKCTCAACACCCTGGT CAAGCAGCTGTCC |
| Cassette10_5 | CTGCAGGACGTGGTCAACCAGAATGCCCAAGCCNNKAACACCCTGGT CAAGCAGCTGTCC |
| Cassette10_6 | CTGCAGGACGTGGTCAACCAGAACGCACAAGCCCTCNNKACCCTGGT CAAGCAGCTGTCC |
| Cassette10_7 | CTGCAGGACGTGGTCAACCAGAACGCACAGGCACTCAATNNKCTGGT CAAGCAGCTGTCC |
| Cassette10_8 | CTGCAGGACGTGGTCAACCAGAACGCCCAAGCACTGAATACCNNKGT CAAGCAGCTGTCC |
| Cassette11_1 | GCCCAGGCACTGAACACCCTGNNKAAACAACTGTCCTCCAACTTCGG CGCCATCAGCTCT |
| Cassette11_2 | GCCCAGGCACTGAACACCCTGGTANNKCAGCTCTCCTCCAACTTCGG CGCCATCAGCTCT |
| Cassette11_3 | GCCCAGGCACTGAACACCCTGGTAAAGNNKCTGTCTTCCAACTTCGG CGCCATCAGCTCT |
| Cassette11_4 | GCCCAGGCACTGAACACCCTGGTCAAACAGNNKTCTTCCAACTTCGG CGCCATCAGCTCT |
| Cassette11_5 | GCCCAGGCACTGAACACCCTGGTCAAGCAACTCNNKTCCAACTTCGG CGCCATCAGCTCT |
| Cassette11_6 | GCCCAGGCACTGAACACCCTGGTAAAACAACTCTCTNNKAACTTCGG CGCCATCAGCTCT |
| Cassette11_7 | GCCCAGGCACTGAACACCCTGGTAAAGCAACTCTCCTCTNNKTTCGG CGCCATCAGCTCT |
| Cassette11_8 | GCCCAGGCACTGAACACCCTGGTAAAGCAGCTCTCTTCTAACNNKGG CGCCATCAGCTCT |
| Cassette12_1 | AAGCAGCTGTCCTCCAACTTCNNKGCAATAAGCTCTGTGCTGAACGAT ATCCTGAGCAGA |
| Cassette12_2 | AAGCAGCTGTCCTCCAACTTCGGTNNKATCAGTTCTGTGCTGAACGAT ATCCTGAGCAGA |
| Cassette12_3 | AAGCAGCTGTCCTCCAACTTCGGTGCCNNKAGCTCGGTGCTGAACGA TATCCTGAGCAGA |
| Cassette12_4 | AAGCAGCTGTCCTCCAACTTCGGCGCAATCNNKTCGGTGCTGAACGA TATCCTGAGCAGA |
| Cassette12_5 | AAGCAGCTGTCCTCCAACTTCGGCGCCATAAGTNNKGTGCTGAACGA TATCCTGAGCAGA |
| Cassette12_6 | AAGCAGCTGTCCTCCAACTTCGGTGCAATAAGTTCGNNKCTGAACGAT ATCCTGAGCAGA |
| Cassette12_7 | AAGCAGCTGTCCTCCAACTTCGGTGCAATCAGCTCTGTANNKAACGAT ATCCTGAGCAGA |
| Cassette12_8 | AAGCAGCTGTCCTCCAACTTCGGTGCCATCAGTTCGGTACTGNNKGAT ATCCTGAGCAGA |
| Cassette13_1 | GCCATCAGCTCTGTGCTGAACNNKATACTCAGCAGACTGGACAAGGT GGAAGCCGAGGTG |
| Cassette13_2 | GCCATCAGCTCTGTGCTGAACGACNNKCTGAGTAGACTGGACAAGGT GGAAGCCGAGGTG |

| | |
|---|---|
| Cassette13_3 | GCCATCAGCTCTGTGCTGAACGATATCNNKAGTAGGCTGGACAAGGTGGAAGCCGAGGTG |
| Cassette13_4 | GCCATCAGCTCTGTGCTGAACGATATCCTCNNKAGACTCGACAAGGTGGAAGCCGAGGTG |
| Cassette13_5 | GCCATCAGCTCTGTGCTGAACGATATACTGAGCNNKCTCGACAAGGTGGAAGCCGAGGTG |
| Cassette13_6 | GCCATCAGCTCTGTGCTGAACGACATCCTCAGCAGGNNKGACAAGGTGGAAGCCGAGGTG |
| Cassette13_7 | GCCATCAGCTCTGTGCTGAACGACATACTCAGTAGGCTGNNKAAGGTGGAAGCCGAGGTG |
| Cassette13_8 | GCCATCAGCTCTGTGCTGAACGACATACTGAGCAGGCTGGACNNKGTGGAAGCCGAGGTG |
| Cassette14_1 | ATCCTGAGCAGACTGGACAAGNNKGAGGCAGAGGTGCAGATCGACAGACTGATCACCGGA |
| Cassette14_2 | ATCCTGAGCAGACTGGACAAGGTANNKGCCGAAGTGCAGATCGACAGACTGATCACCGGA |
| Cassette14_3 | ATCCTGAGCAGACTGGACAAGGTAGAANNKGAGGTACAGATCGACAGACTGATCACCGGA |
| Cassette14_4 | ATCCTGAGCAGACTGGACAAGGTGGAGGCCNNKGTACAGATCGACAGACTGATCACCGGA |
| Cassette14_5 | ATCCTGAGCAGACTGGACAAGGTGGAAGCAGAANNKCAGATCGACAGACTGATCACCGGA |
| Cassette14_6 | ATCCTGAGCAGACTGGACAAGGTAGAGGCAGAAGTANNKATCGACAGACTGATCACCGGA |
| Cassette14_7 | ATCCTGAGCAGACTGGACAAGGTAGAGGCCGAGGTACAANNKGACAGACTGATCACCGGA |
| Cassette14_8 | ATCCTGAGCAGACTGGACAAGGTAGAAGCAGAGGTGCAAATCNNKAGACTGATCACCGGA |
| Cassette15_1 | GAAGCCGAGGTGCAGATCGACNNKCTCATAACCGGAAGGCTGCAGTCCCTGCAGACCTAC |
| Cassette15_2 | GAAGCCGAGGTGCAGATCGACAGGNNKATCACAGGAAGGCTGCAGTCCCTGCAGACCTAC |
| Cassette15_3 | GAAGCCGAGGTGCAGATCGACAGGCTGNNKACCGGTAGGCTGCAGTCCCTGCAGACCTAC |
| Cassette15_4 | GAAGCCGAGGTGCAGATCGACAGACTCATCNNKGGTAGGCTGCAGTCCCTGCAGACCTAC |
| Cassette15_5 | GAAGCCGAGGTGCAGATCGACAGACTGATAACANNKAGGCTGCAGTCCCTGCAGACCTAC |
| Cassette15_6 | GAAGCCGAGGTGCAGATCGACAGGCTCATAACAGGTNNKCTGCAGTCCCTGCAGACCTAC |
| Cassette15_7 | GAAGCCGAGGTGCAGATCGACAGGCTCATCACCGGTAGANNKCAGTCCCTGCAGACCTAC |
| Cassette15_8 | GAAGCCGAGGTGCAGATCGACAGGCTGATCACCGGAAGACTGNNKTCCCTGCAGACCTAC |
| Cassette16_1 | CTGATCACCGGAAGGCTGCAGNNKCTCCAAACCTACGTTACCCAGCAGCTGATCAGAGCC |
| Cassette16_2 | CTGATCACCGGAAGGCTGCAGTCTNNKCAGACATACGTTACCCAGCAGCTGATCAGAGCC |
| Cassette16_3 | CTGATCACCGGAAGGCTGCAGTCTCTGNNKACCTATGTTACCCAGCAGCTGATCAGAGCC |

| | |
|---|---|
| Cassette16_4 | CTGATCACCGGAAGGCTGCAGTCCCTCCAGNNKTATGTTACCCAGCA GCTGATCAGAGCC |
| Cassette16_5 | CTGATCACCGGAAGGCTGCAGTCCCTGCAAACANNKGTTACCCAGCA GCTGATCAGAGCC |
| Cassette16_6 | CTGATCACCGGAAGGCTGCAGTCTCTCCAAACATATNNKACCCAGCA GCTGATCAGAGCC |
| Cassette16_7 | CTGATCACCGGAAGGCTGCAGTCTCTCCAGACCTATGTANNKCAGCA GCTGATCAGAGCC |
| Cassette16_8 | CTGATCACCGGAAGGCTGCAGTCTCTGCAAACCTACGTAACCNNKCA GCTGATCAGAGCC |
| Cassette17_1 | CTGCAGACCTACGTTACCCAGNNKCTCATAAGAGCCGCCGAGATTAG AGCCTCTGCCAAT |
| Cassette17_2 | CTGCAGACCTACGTTACCCAGCAANNKATCAGGGCCGCCGAGATTAG AGCCTCTGCCAAT |
| Cassette17_3 | CTGCAGACCTACGTTACCCAGCAACTGNNKAGAGCAGCCGAGATTAG AGCCTCTGCCAAT |
| Cassette17_4 | CTGCAGACCTACGTTACCCAGCAGCTCATCNNKGCAGCCGAGATTAG AGCCTCTGCCAAT |
| Cassette17_5 | CTGCAGACCTACGTTACCCAGCAGCTGATAAGGNNKGCCGAGATTAG AGCCTCTGCCAAT |
| Cassette17_6 | CTGCAGACCTACGTTACCCAGCAACTCATAAGGGCANNKGAGATTAG AGCCTCTGCCAAT |
| Cassette17_7 | CTGCAGACCTACGTTACCCAGCAACTCATCAGAGCAGCANNKATTAGA GCCTCTGCCAAT |
| Cassette17_8 | CTGCAGACCTACGTTACCCAGCAACTGATCAGGGCAGCAGAGNNKAG AGCCTCTGCCAAT |
| Cassette18_1 | CTGATCAGAGCCGCCGAGATTNNKGCATCGGCCAATCTGGCCGCCAC CAAGATGTCTGAG |
| Cassette18_2 | CTGATCAGAGCCGCCGAGATTAGGNNKTCTGCAAATCTGGCCGCCAC CAAGATGTCTGAG |
| Cassette18_3 | CTGATCAGAGCCGCCGAGATTAGGGCCNNKGCCAACCTGGCCGCCA CCAAGATGTCTGAG |
| Cassette18_4 | CTGATCAGAGCCGCCGAGATTAGAGCATCTNNKAACCTGGCCGCCAC CAAGATGTCTGAG |
| Cassette18_5 | CTGATCAGAGCCGCCGAGATTAGAGCCTCGGCANNKCTGGCCGCCA CCAAGATGTCTGAG |
| Cassette18_6 | CTGATCAGAGCCGCCGAGATTAGGGCATCGGCAAACNNKGCCGCCA CCAAGATGTCTGAG |
| Cassette18_7 | CTGATCAGAGCCGCCGAGATTAGGGCATCTGCCAACCTCNNKGCCAC CAAGATGTCTGAG |
| Cassette18_8 | CTGATCAGAGCCGCCGAGATTAGGGCCTCTGCAAACCTCGCCNNKAC CAAGATGTCTGAG |
| Cassette19_1 | GCCTCTGCCAATCTGGCCGCCNNKAAAATGTCGGAGTGTGTGCTGGG CCAGAGCAAGAGA |
| Cassette19_2 | GCCTCTGCCAATCTGGCCGCCACANNKATGTCTGAATGTGTGCTGGG CCAGAGCAAGAGA |
| Cassette19_3 | GCCTCTGCCAATCTGGCCGCCACCAAGNNKTCGGAATGTGTGCTGGG CCAGAGCAAGAGA |
| Cassette19_4 | GCCTCTGCCAATCTGGCCGCCACCAAAATGNNKGAATGCGTGCTGGG CCAGAGCAAGAGA |

| | |
|---|---|
| Cassette19_5 | GCCTCTGCCAATCTGGCCGCCACAAAGATGTCGNNKTGCGTGCTGGGCCAGAGCAAGAGA |
| Cassette19_6 | GCCTCTGCCAATCTGGCCGCCACCAAGATGTCTGAANNKGTACTGGGCCAGAGCAAGAGA |
| Cassette19_7 | GCCTCTGCCAATCTGGCCGCCACAAAAATGTCTGAGTGCNNKCTGGGCCAGAGCAAGAGA |
| Cassette19_8 | GCCTCTGCCAATCTGGCCGCCACCAAGATGTCGGAGTGTGTANNKGGCCAGAGCAAGAGA |

| | |
|---|---|
| Cassette1_Rprimer | GATTGTGCCGGCCAGCAGGGC |
| Cassette2_Rprimer | AGCTCCAAATGTCCAGCCGCT |
| Cassette3_Rprimer | AAAGGGGATCTGCAGAGCGGC |
| Cassette4_Rprimer | GAACCGGTAGGCCATCTGCAT |
| Cassette5_Rprimer | ATTCTGGGTCACTCCGATGCC |
| Cassette6_Rprimer | CAGCTTCTGGTTCTCGTACAG |
| Cassette7_Rprimer | GGCGCTGTTGAACTGGTTGGC |
| Cassette8_Rprimer | CAGGCTGTCCTGGATCTTGCC |
| Cassette9_Rprimer | TCCCAGGGCGCTTGCTGTGCT |
| Cassette10_Rprimer | CTGGTTGACCACGTCCTGCAG |
| Cassette11_Rprimer | CAGGGTGTTCAGTGCCTGGGC |
| Cassette12_Rprimer | GAAGTTGGAGGACAGCTGCTT |
| Cassette13_Rprimer | GTTCAGCACAGAGCTGATGGC |
| Cassette14_Rprimer | CTTGTCCAGTCTGCTCAGGAT |
| Cassette15_Rprimer | GTCGATCTGCACCTCGGCTTC |
| Cassette16_Rprimer | CTGCAGCCTTCCGGTGATCAG |
| Cassette17_Rprimer | CTGGGTAACGTAGGTCTGCAG |
| Cassette18_Rprimer | AATCTCGGCGGCTCTGATCAG |
| Cassette19_Rprimer | GGCGGCCAGATTGGCAGAGGC |

33

**Supplementary Table 4.** Numbers of cells collected per bin in expression sorting.

| Bin | Replicate 1 | Replicate 2 | Replicate 3 |
|---|---|---|---|
| Bin 0 | $8.03 \times 10^5$ | $1.30 \times 10^6$ | $1.70 \times 10^6$ |
| Bin 1 | $7.51 \times 10^5$ | $1.30 \times 10^6$ | $1.70 \times 10^6$ |
| Bin 2 | $7.70 \times 10^5$ | $1.30 \times 10^6$ | $1.70 \times 10^6$ |
| Bin 3 | $8.20 \times 10^5$ | $1.30 \times 10^6$ | $1.70 \times 10^6$ |

**Supplementary Table 5.** Numbers of cells collected per bin in fusion sorting.

| Bin | Replicate 1 | Replicate 2 |
|---|---|---|
| $mNG2^-$ | $3.53 \times 10^6$ | $5.51 \times 10^6$ |
| $mNG2^+$ | $1.84 \times 10^5$ | $4.87 \times 10^5$ |

**Supplementary Table 6.** *p*-values from Student's *t* test of expression and fusion scores between mutation types. Data are related to **Extended Data Fig. 2c,d**.

| **Expression** | | | |
|---|---|---|---|
| | Missense | Nonsense | Silent |
| Missense | | $6.46 \times 10^{-60}$ | $2.93 \times 10^{-2}$ |
| Nonsense | $6.46 \times 10^{-60}$ | | $3.80 \times 10^{-44}$ |
| Silent | $2.93 \times 10^{-2}$ | $3.80 \times 10^{-44}$ | |
| **Fusion** | | | |
| | Missense | Nonsense | Silent |
| Missense | | $3.10 \times 10^{-34}$ | $7.08 \times 10^{-4}$ |
| Nonsense | $3.10 \times 10^{-34}$ | | $1.44 \times 10^{-31}$ |
| Silent | $7.08 \times 10^{-4}$ | $1.44 \times 10^{-31}$ | |

**Supplementary Table 7.** Betacoronaviruses used for sequence conservation analysis. Data are related to **Extended Data Fig. 3c,d**.

| Accession ID | Database | Name |
|---|---|---|
| gb_MN908947.3_ | GenBank | Severe acute respiratory syndrome coronavirus 2 isolate Wuhan-Hu-1, complete genome |
| gb_MN996532.2_ | GenBank | Bat coronavirus RaTG13, complete genome |
| gb_MZ937000.1_ | GenBank | Bat coronavirus isolate BANAL-20-52/Laos/2020, complete genome |
| EPI_ISL_410543 | GISAID | hCoV-19/pangolin/Guangxi/P3B/2017 |
| EPI_ISL_471465 | GISAID | hCoV-19/pangolin/Guangdong/cDNA20-S/2019 |
| gb_MZ937003.1_ | GenBank | Bat coronavirus isolate BANAL-20-236/Laos/2020, complete genome |
| gb_MZ937001.1_ | GenBank | Bat coronavirus isolate BANAL-20-103/Laos/2020, complete genome |
| gb_MG772933.1_ | GenBank | Bat SARS-like coronavirus isolate bat-SL-CoVZC45, complete genome |
| gb_MG772934.1_ | GenBank | Bat SARS-like coronavirus isolate bat-SL-CoVZXC21, complete genome |
| gb_KT444582.1_ | GenBank | SARS-like coronavirus WIV16, complete genome |
| gb_DQ497008.1_ | GenBank | SARS coronavirus strain MA-15, complete genome |
| gb_KC881007.1_ | GenBank | Bat SARS-like coronavirus WIV1 spike protein (S) gene, complete cds |
| gb_KY417144.1_ | GenBank | Bat SARS-like coronavirus isolate Rs4084, complete genome |
| gb_DQ412042.1_ | GenBank | Bat SARS coronavirus Rf1, complete genome |
| gb_KJ473813.1_ | GenBank | BtRf-BetaCoV/SX2013, complete genome |
| gb_KJ473815.1_ | GenBank | BtRs-BetaCoV/GX2013, complete genome |
| gb_KF294457.1_ | GenBank | Bat SARS-like coronavirus isolate Longquan-140 orf1ab polyprotein, spike glycoprotein, envelope protein, membrane protein, and nucleocapsid protein genes, complete cds |
| gb_DQ022305.2_ | GenBank | Bat SARS coronavirus HKU3-1, complete genome |
| gb_DQ071615.1_ | GenBank | Bat SARS coronavirus Rp3, complete genome |
| gb_FJ588686.1_ | GenBank | Bat SARS CoV Rs672/2006, complete genome |
| gb_KJ473814.1_ | GenBank | BtRs-BetaCoV/HuB2013, complete genome |
| gb_KF569996.1_ | GenBank | *Rhinolophus affinis* coronavirus isolate LYRa11, complete genome |
| gb_KY352407.1_ | GenBank | Severe acute respiratory syndrome-related coronavirus strain BtKY72, complete genome |
| ref_NC_014470.1_ | GenBank | Bat coronavirus BM48-31/BGR/2008, complete genome |
| gb_MZ937004.1_ | GenBank | Bat coronavirus isolate BANAL-20-247/Laos/2020, complete genome |
| gb_MZ937002.1_ | GenBank | Bat coronavirus isolate BANAL-20-116/Laos/2020, complete genome |
| EPI_ISL_412977 | GISAID | hCoV-19/bat/Yunnan/RmYN02/2019 |
